# Hyperglycemia Activates Retinal Photoreceptors to Induce Neuroglial Inflammation

**DOI:** 10.64898/2026.08.27.747567

**Authors:** Jorge L. Nunez, Jacob M. Poloway, Yanxin Chen, Devyn H. Zaminski, John S. Penn, MD Imam Uddin, Irina De la Huerta

**Affiliations:** Vanderbilt University Medical Center, Nashville, TN, USA; Vanderbilt University, Nashville, TN, USA; Department of Ophthalmology and Visual Sciences, Vanderbilt University School of Medicine, Nashville, TN, USA

**Author notes:** Corresponding author: Irina De la Huerta, MD, PhD, Department of Ophthalmology and Visual Sciences Vanderbilt University School of Medicine, 2311 Pierce Avenue, Nashville, TN, 37232-8808. Disclosures: **J.L. Nunez**, None; **J.M. Poloway**, None; **Y. Chen**, None; **D.H. Zaminsky**, None; **J.S. Penn**, None; **M.I. Uddin**, None. **I. De la Huerta**, None.

## Abstract

Diabetic retinopathy (DR) is a major cause of vision loss in working-age adults. Accumulating evidence suggests that retinal photoreceptors contribute to the initiation and progression of diabetic retinopathy. In this study, we investigated whether hyperglycemia directly alters photoreceptor signaling and whether photoreceptor-derived inflammatory mediators activate downstream Müller glial cells. Primary photoreceptors were isolated from adult mice and cultured with normal glucose, high D-glucose, or high L-glucose as an osmotic control. Photoreceptor-conditioned media were analyzed for inflammatory and growth factors and used to stimulate primary Müller glia. High glucose exposure increased photoreceptor production of TNF-α, IL-6, and VEGF. Photoreceptor-conditioned media from high glucose-treated photoreceptors induced Müller glial expression of IL-1β, TNF-α, and IL-6. Müller glia exposed to photoreceptor-conditioned media increased VEGF expression and secretion and enhanced MMP-9 expression, secretion, and gelatinase activity. Together, these findings support a direct role for photoreceptors as glucose-responsive neuronal cells that can initiate and amplify neuroglial inflammatory signaling in hyperglycemic conditions.

## Introduction

Diabetic retinopathy (DR) is a leading cause of vision loss in working-age adults (https://usdss.cdc.gov/diabetes/report.html).^1–4^ Although vascular abnormalities are prominent clinical features of DR, neuronal and glial dysfunction develops early and contributes to retinal pathology.^5–9^ Inflammatory cytokines and growth factors, including TNF-α, IL-6, IL-1β, and VEGF, are increased in diabetic retinas and are associated with retinal inflammation and disease progression.^10–12^ The cellular sources and upstream signals responsible for these changes, particularly before advanced vascular pathology develops, remain incompletely understood.

Hypoxia has been implicated as a cause of increased VEGF and inflammation in DR.^13, 14^ Hypoxic Müller glial cells are the predominant source of VEGF in the ischemic retina, and inhibiting Müller glial VEGF production reduces inflammatory signaling.^15, 16^ However, retinal hypoxia is not present in early diabetes, at the time when inflammatory mediators and VEGF levels become significantly increased in the diabetic retina.^17^ These observations raise the possibility that glucose-sensitive neural cells contribute to early inflammatory signaling independently of hypoxia.

Accumulating evidence suggests that rod photoreceptors may play a role in the development and progression of diabetic retinopathy.^18–22^ Photoreceptor ablation, whether genetic or pharmacologic, decreases levels of inflammatory mediators in the retinas of diabetic mice.^18–20^ Müller glial cells are closely associated with photoreceptors and respond to extracellular signals released by retinal neurons.^23, 24^ These findings suggest that photoreceptors may function as upstream sensors of metabolic stress and that Müller glia may amplify signals initiated by photoreceptors.

In the present study, we used primary retinal photoreceptors to investigate whether physiologically relevant high glucose levels directly stimulate photoreceptors to produce inflammatory mediators and whether photoreceptor-derived inflammatory mediators could activate Müller glia. We provide evidence that hyperglycemia increases photoreceptor production of TNF-α, IL-6, and VEGF and that conditioned media from high glucose photoreceptors stimulate Müller glial production of VEGF and MMP-9. These findings support a photoreceptorto-Müller glia signaling pathway that may contribute to neuroglial inflammation in the diabetic retina (**Figure 1**).

**Figure 1.**
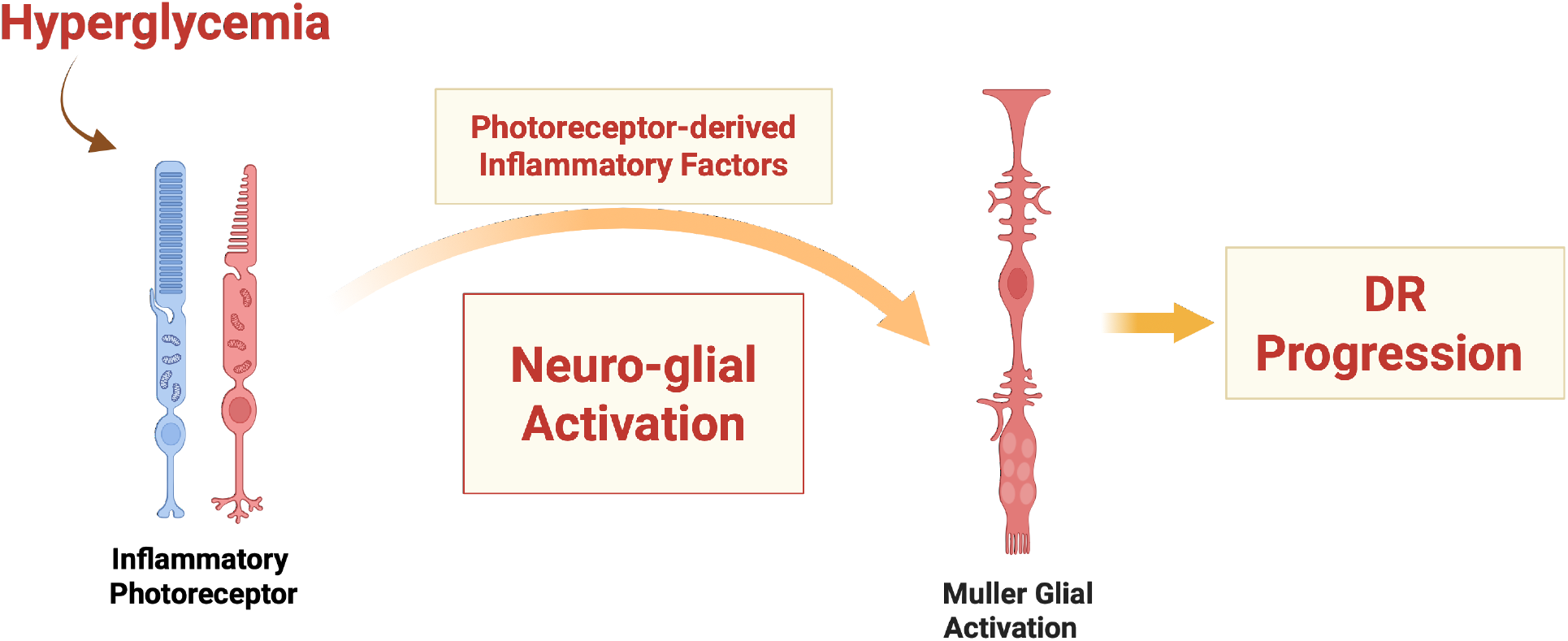
Hyperglycemia causes neuroglial activation driven by inflammation. In diabetic retina hyperglycemia triggers photoreceptor activation, a process that contributes to DR progression. Hyperglycemia initiates photoreceptor activation causing release of inflammatory factors that in turn activate Müller glia, leading to DR progression.

## Materials and Methods

### Animals

All experiments were approved by the Vanderbilt University Institutional Animal Care and Use Committee and were performed in accordance with the ARVO Statement for the Use of Animals in Ophthalmic and Vision Research. Eight C57BL/6 mice aged 3 weeks old (Charles River, Wilmington, MA) were used for each cell culture isolation. Equal numbers of males and female mice were used.

### Cell culture

1. Primary rod photoreceptor cultures were generated using previously published protocols adapted by us for culturing mature mouse photoreceptors.^25–27^ Retinas were dissected from C57BL/6J mice in Hank’s Balanced Salt Solution (HBSS) (GIBCO) on ice and enzymatically dissociated using the Worthington papain dissociation system (Worthington). Retinas were washed again with 0.5% BSA in AMES buffer and retinal tissue was triturated 6X with a P1000 pipette tip. Retinal cells were suspended in 300 µl 0.5% BSA in AMES and incubated with 2 µl rabbit anti-CD73 (Abcam cat#175396) per 10^7^ cells for 15 minutes (min) at room temperature. Cells were centrifuged, resuspended in fresh 0.5% BSA in AMES, and incubated with 20 µl goat anti-rabbit magnetic microbeads (Miltenyi Biotec cat#130-048-602) for 15 min at room temperature. CD73-positive cells were separated from unbound cells in a magnetic column. CD73-positive cells were seeded at 200,000 cells/well on glass coverslips coated with poly-D-ornithine in 24 well plates. Cells were cultured in Dulbecco’s Modified Eagle Medium (DMEM) containing 1% N1 supplement,1X penicillin-streptomycin (Sigma), and either 5 mM D-glucose, or 30 mM D-glucose, or 5 mM D-glucose + 25 mM L-glucose as osmotic control. Rod photoreceptor enrichment to at least 98% was confirmed by immunostaining for rhodopsin.
2. Primary mouse Müller glia were isolated from C57BL/6 retinas that were dissected from C57BL/6J mice in Hank’s Balanced Salt Solution (HBSS) (GIBCO) on ice and enzymatically dissociated using collagenase and trypsin. Retinas were washed again with 0.5% BSA in AMES buffer and retinal tissue was triturated 6X with a P1000 pipette tip. Retinal cells were suspended in 300 µl 0.5% BSA in AMES and incubated with 2 µl rabbit anticellular retinaldehyde-binding protein (CRALBP) (Invitrogen cat# MA55033) per 10^7^ cells for 15 minutes (min) at room temperature. Cells were centrifuged, resuspended in fresh 0.5% BSA in AMES, and incubated with 20 µl goat anti-rabbit magnetic microbeads (Miltenyi Biotec cat#130-048-602) for 15 min at room temperature. CRALBP-positive cells were separated from unbound cells in a magnetic column and cultured in DMEM with 10% FBS. Müller cell identity was confirmed by labeling with antibodies to glial fibrillary acid protein (GFAP) and to CRALBP. Müller glia from passages 4-8 were used for experiments.

### Immunohistochemistry

Retinal cells on coverslips were fixed in 4% paraformaldehyde for 15 min, washed with PBS and incubated in blocking buffer (0.3% Triton X-100 and 3% goat serum in PBS) for 30 min at room temperature. Coverslips were incubated with primary antibodies overnight at 4°C using anti-rhodopsin (1:150), anti-GFAP (1:400), and anti-CRALBP (1:500). Slides were washed 3 times in PBS and incubated with Alexa Fluor secondary antibodies (Invitrogen) diluted 1:1000 for 2 h at room temperature. Stained coverslips were washed 3 times in PBS and mounted in DAPI Fluoromount-G.

### Enzyme-linked immunosorbent assay (ELISA)

Primary photoreceptor cultures were treated with 5 mM D-glucose or 30 mM D-glucose, or 5 mM D-glucose + 25 mM L-glucose as osmotic control for 48 hours (h). The cell culture supernatant was concentrated two-fold using 3kDa concentrators (cat# 28932240, Cytiva, UK) and processed using mouse high sensitivity ELISA kits specific for VEGF (cat# EMVEGFACL), and for cytokines IL-6 (cat# BMS603HS), IL-8 (cat# EMCXCL15), IL-1β (cat# BMS6002), and TNF-α (cat# BMS607HS). All ELISA kits were purchased from Invitrogen; Waltham, MA. The minimum detection limits were: VEGF 2 pg/ml, TNF-α 0.75 pg/ml, IL-6 0.21 pg/ml, IL-1β 1.2 pg/ml, IL-8 0.8 pg/ml.

### RNA Isolation and Quantitative Real Time (qRT) PCR

Cultured Müller glia were lysed and total RNA was extracted using an RNeasy Mini kit (Qiagen, Valencia, CA). The High-Capacity cDNA Archive Kit (Applied Biosystems, Waltham, MA) was used to reverse transcribe the total RNA. qRT-PCR was performed using TaqMan Gene Expression Assays (Applied Biosystems). The coamplification of TNF-α, IL-6, IL-1β, VEGF, and MMP-9 was compared with 18S as the normalization control. Data were analyzed using the comparative Ct method. A minimum of 4 independent samples per group and technical replicates of each sample were used in all experiments.

### Western blot and zymography

Cell lysates and conditioned media were run on 4-15% gradient precast polyacrylamide gels (Bio-Rad). Revert 700 total protein stain (LI-COR) was used post-transfer for total protein quantification and normalization analysis. Immunoblotting was performed using rabbit monoclonal antibodies to VEGF (Abcam cat#ab51745) or MMP2 (Abcam cat#ab92536) or MMP9 (Abcam cat#38898). Gelatinase enzymatic activity was measured by zymographic assays using Novex Zymogram Plus (10% Gelatin) gels (Invitrogen).

### Statistical analysis

Data were analyzed using Prism software (GraphPad; La Jolla, CA). T-test and ANOVA with Tukey’s multiple comparisons post-hoc test were used to evaluate significant differences among treatment groups. Values of p < 0.05 were considered statistically significant.

## RESULTS

### Effect of elevated glucose levels on photoreceptor production of inflammatory mediators

To study the effect of high glucose on photoreceptors in vitro, we developed a protocol for isolating adult mouse rod photoreceptors and used it to generate primary dissociated photoreceptor cultures. We incubated photoreceptors for 48 h with physiologically relevant high D-glucose levels (30 mM D-glucose), with physiologically normal D-glucose levels (5 mM D-glucose), or with high L-glucose as an osmotic control (5 mM D-glucose + 25 mM L-glucose). We measured the concentrations of cytokines and growth factors in the photoreceptor conditioned media via multiplex ELISA. Media from photoreceptors treated with elevated D-glucose contained significantly higher concentrations of TNF-α, VEGF, and IL-6 compared with media from photoreceptors treated with normal D-glucose or with L-glucose as an osmotic control (**Figure 2**). Other inflammatory cytokines such as IL-1β and IL-8 (CXCL15) were not detected.

**Figure 2.**
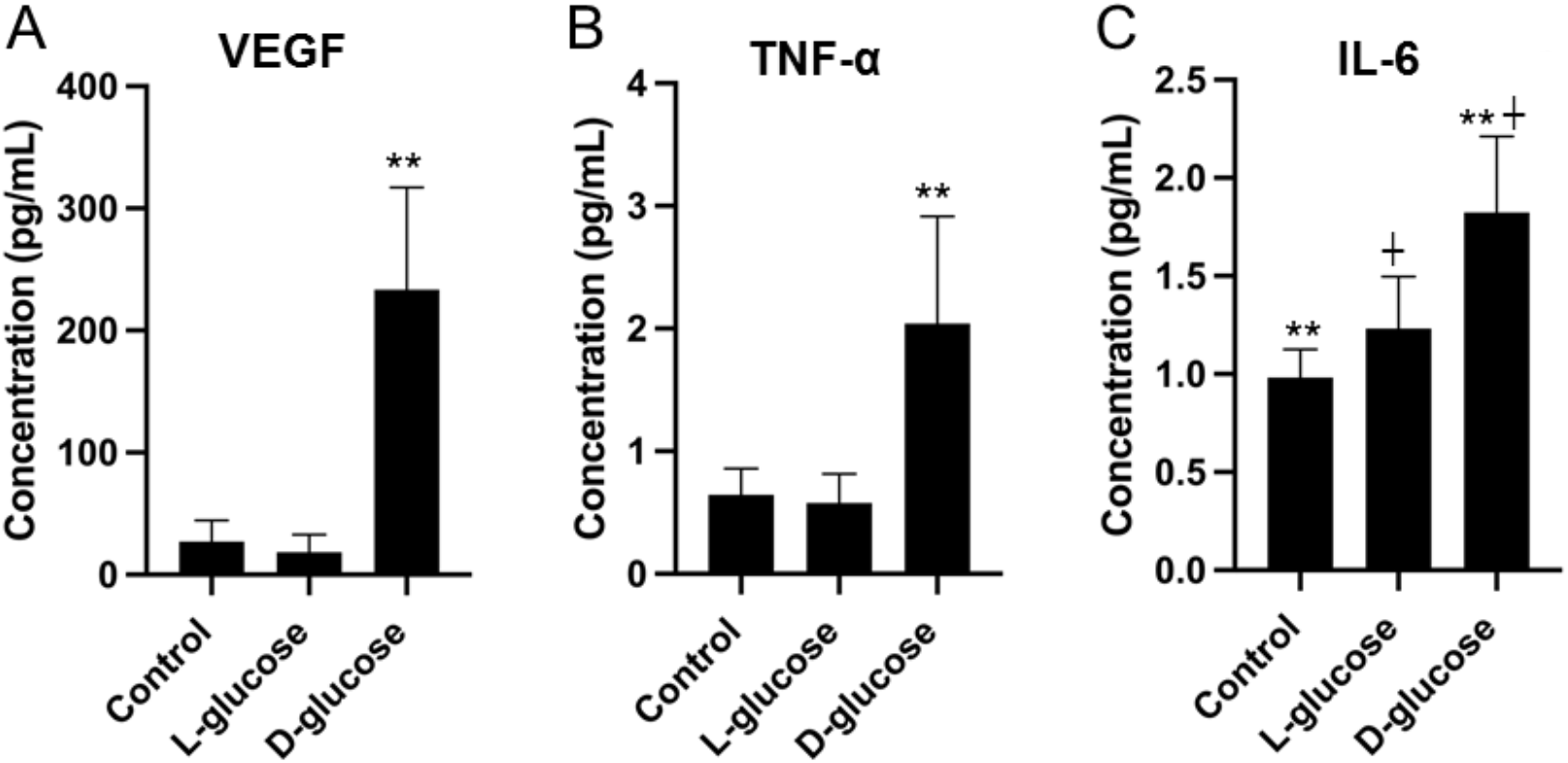
Photoreceptors release pro-angiogenic and pro-inflammatory mediators when exposed to elevated glucose levels. ELISA assay for (A) VEGF, (B) TNF-α, and (C) IL-6 concentrations in medium from rod photoreceptors treated with high D-glucose compared to normal D-glucose and to high L-glucose (osmotic control). Normal glucose = 5 mM D-glucose, high D-glucose = 30 mM D-glucose, high L-glucose = 5 mM D-glucose plus 25 mM L-glucose. Results are expressed as the mean ± standard deviation (n=6). **p < 0.01 high D-glucose vs. control and high D-glucose vs. high L-glucose. †p<0.05 high D-glucose vs. high L-glucose.

### Photoreceptors exposed to high glucose induce Müller glial cell expression of inflammatory cytokines Il-1β, TNF-α, and IL-6

We investigated the effect of photoreceptor-produced mediators on downstream Müller glial cells. Müller cells are the principal glial cells of the retina and exist in close proximity to photoreceptors, where they process and respond to signals released by retinal neurons. To determine how photoreceptors exposed to elevated glucose levels affect Müller glia we incubated the photoreceptor culture with high glucose for 48 hours. The photoreceptor-conditioned media were then added to Müller glial cells, and the Müller glia were collected 12 hours later for gene expression analysis (Figure 3). The photoreceptor-conditioned media from photoreceptors incubated in high glucose caused a significant induction of *TNF-α, IL-1β* and *IL-*6 expression in Müller glia (p<0.01) (**Figure 3**). These results indicate that under hyperglycemic conditions, photoreceptors can induce a rise in Müller glial expression of inflammatory mediators.

**Figure 3.**
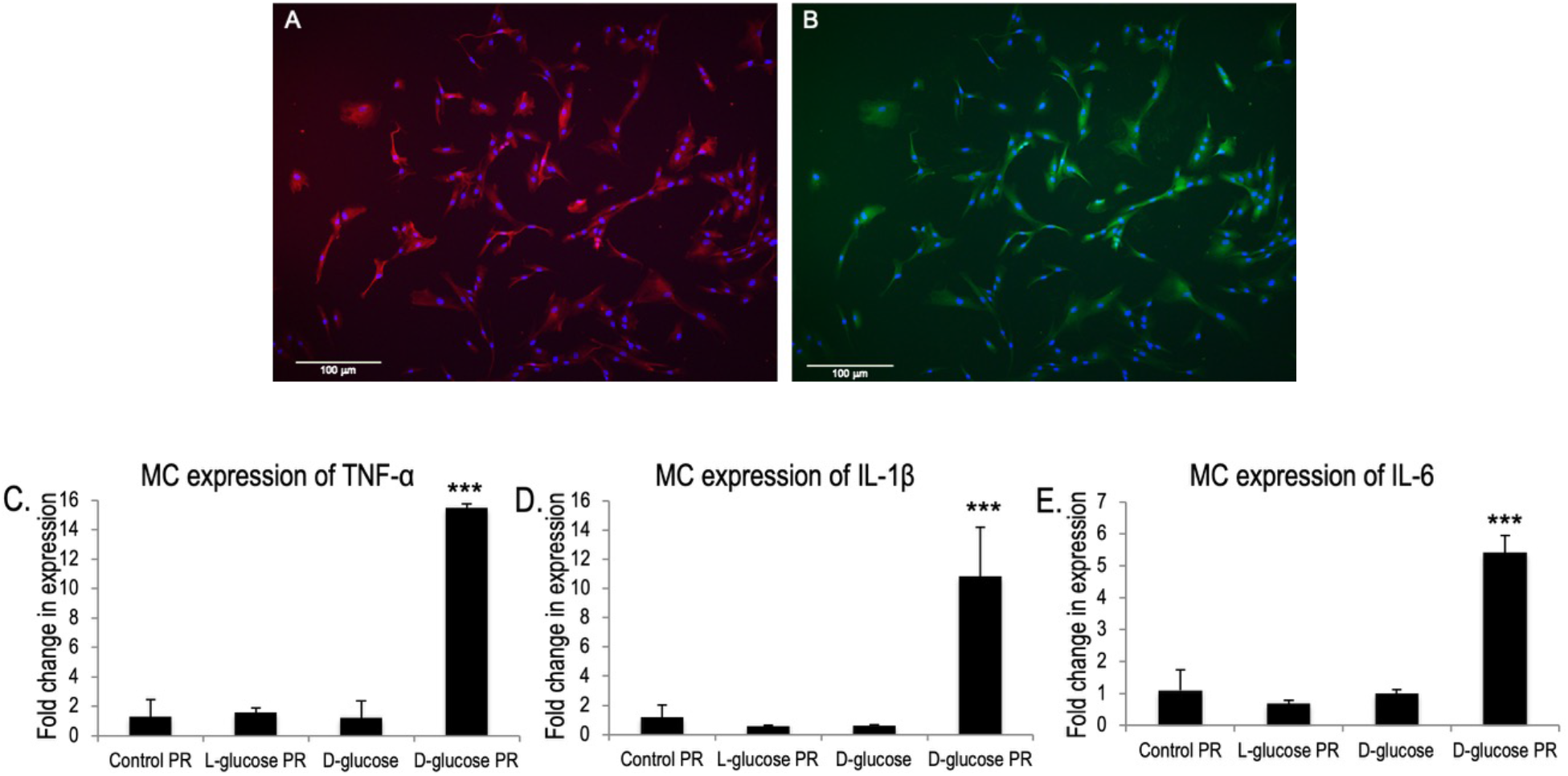
Primary mouse Müller glia culture immunostained for A: CRALBP (red) and DAPI (blue) and B: GFAP (green) and DAPI (blue) to confirm Müller glial cell identity. Scale bar: 100 μm. C-E: Fold changes in TNF-α, IL-1β, and IL-6 expression in Müller glia treated with conditioned media from photoreceptors exposed to normal glucose (Control PR) or from photoreceptors exposed to L-glucose as osmotic control (L-glucose PR) or Müller glia treated with high D-glucose (D-glucose) or with media from photoreceptors incubated with high D-glucose (D-glucose PR). ***p<0.001 vs Control PR. N=6/group.

### Photoreceptors treated with high glucose increase Müller glial production of VEGF

Müller glia are the main producers of VEGF in the diabetic retina.^23, 24^ Müller cells exposed to photoreceptor-conditioned media from photoreceptors treated with high D-glucose had increased expression of VEGF compared with cells treated with conditioned media from photoreceptors exposed to normal D-glucose or high L-glucose as an osmotic control (**Figure 4**). High glucose photoreceptor-conditioned media induced a highly significant 4-fold increase in VEGF secretion by Müller cells compared with Müller cells exposed to high glucose alone.

**Figure 4.**
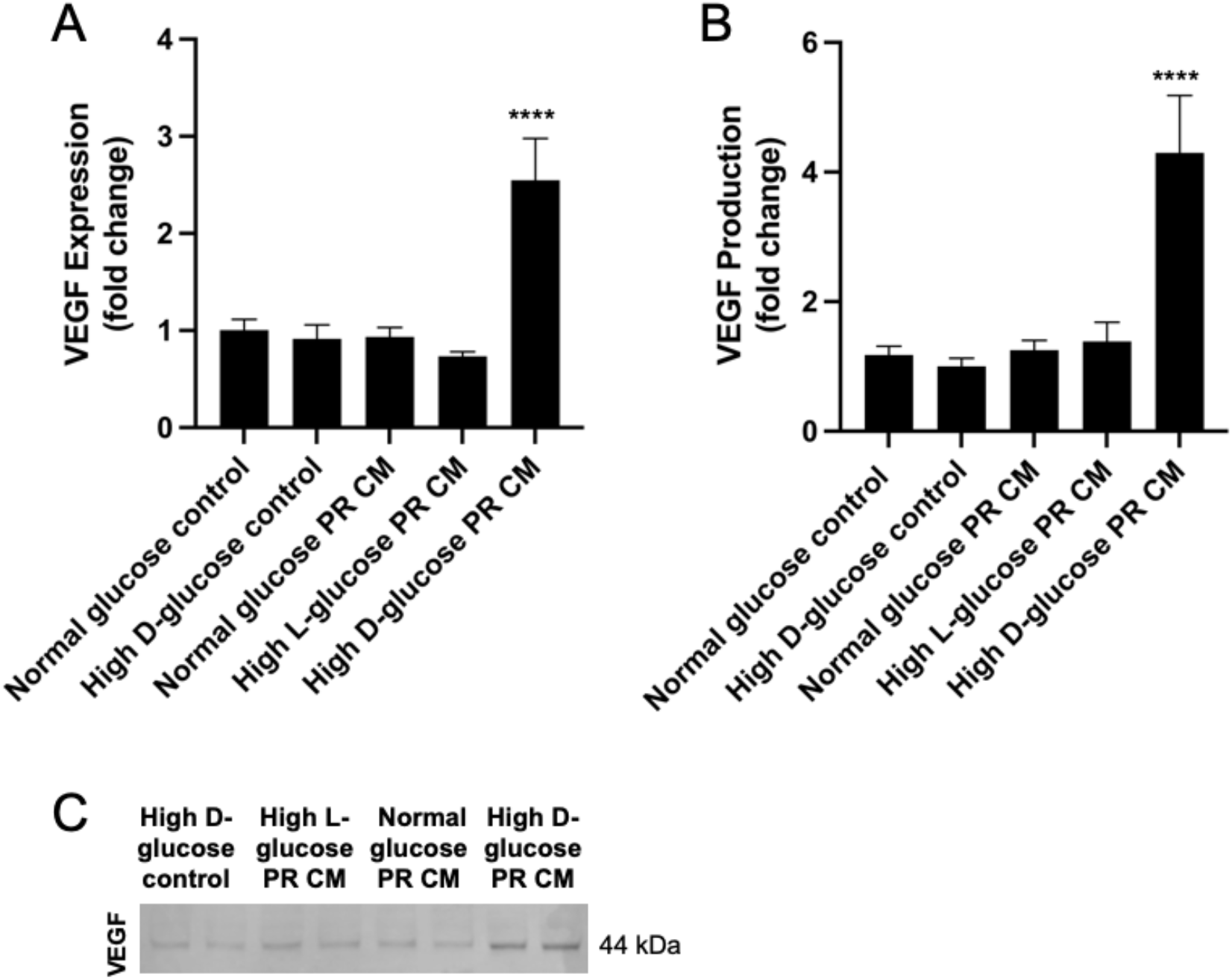
Culture media from photoreceptors exposed to elevated glucose induce Müller cell VEGF expression and production. (A) qRT-PCR for mouse Müller cell VEGF expression after incubation with conditioned media from photoreceptors treated with high D-glucose (High D-glucose PR CM) or with control media. (B, C) Western blot for VEGF secreted by mMC treated with photoreceptor-produced mediators in high glucose (High D-glucose PR CM). Controls include mMC treated with mediators from photoreceptors incubated in normal glucose (Normal glucose PR CM), or in high L-glucose as osmotic control (High L-glucose PR CM). Other controls include mMC treated with normal glucose control medium (Normal glucose control), or with high D-glucose medium (High D-glucose control). VEGF A is secreted as a dimer showing a molecular weight of approximately 44 kDa on immunoblots. Each sample is normalized to total protein. Graphs are normalized to high D-glucose control. Bands are independent duplicates. ****P < 0.0001, n=4.

### Photoreceptors exposed to elevated glucose levels induce Müller cell production of matrix-remodeling mediators

Müller cell expression and secretion of matrix metalloproteinase (MMP)-9 also increased after stimulation with conditioned media from photoreceptors treated with high D-glucose relative to stimulation with media from photoreceptors treated with low D-glucose (**Figure 5**). MMP-9 is an extracellular matrix-remodeling enzyme that has been associated with inflammation and advanced diabetic retinal pathology.^28–30^. To assess MMP enzymatic activity, we performed zymography and found increased Müller cell gelatinase activity after exposure to conditioned media from photoreceptors treated with high D-glucose (**Figure 5**).

**Figure 5.**
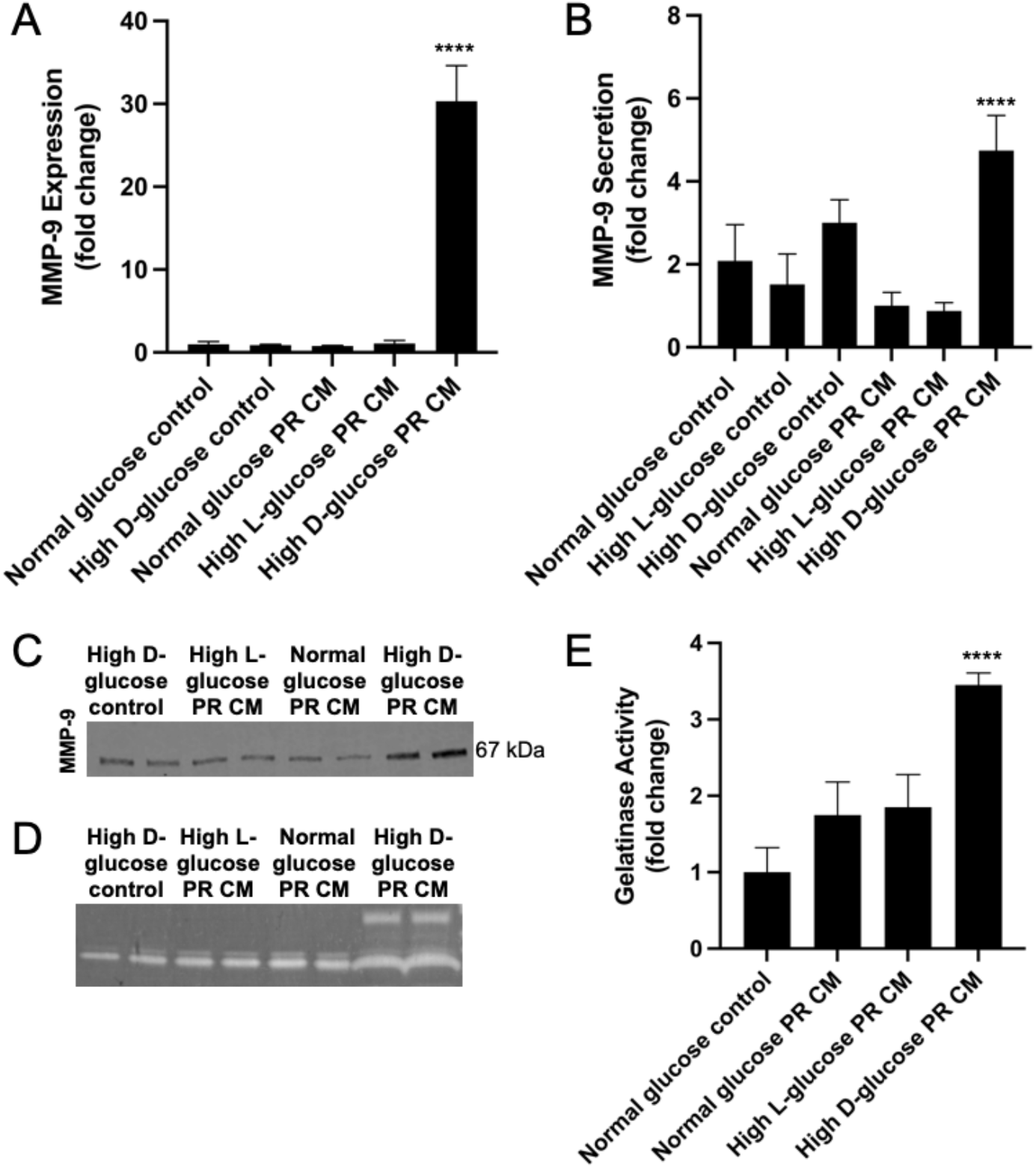
Photoreceptor-produced mediators induce Müller cell expression and production of matrix metalloproteinase 9. (A) qRT-PCR for mouse Müller cell *mmp-9* expression after incubation with conditioned medium from photoreceptors treated with high D-glucose (High D-glucose PR CM) or with control media. Results are reported as fold induction over normal glucose control media. (B, C) Western blot for MMP9 produced by mMC treated with photoreceptor-produced mediators in high glucose (High D-glucose PR CM) or with control media. Each sample is normalized to total protein. Graphs are normalized to Normal glucose PR CM. Bands are independent duplicates. (D, E) Zymogram for mMC-produced gelatinase activity following incubation with conditioned media from photoreceptors treated with high D-glucose (High D-glucose PR CM). Samples are normalized to collagenase control. Graph is normalized to normal glucose control. Bands are independent duplicates. Controls in A-E include mMC treated with media from photoreceptors incubated with normal glucose (Normal glucose PR CM) Normal glucose PR CM, or with media from photoreceptors incubated with high L-glucose as osmotic control (High L-glucose PR CM), or with control media containing high D-glucose (High D-glucose control), or control media containing normal glucose (Normal glucose control). ****P < 0.0001, n=4.

## DISCUSSION

It has been recognized for many years that retinal neuronal damage occurs early in diabetes.^31^ Examination of human post-mortem retinal tissue demonstrated neuronal loss accompanied by increased TUNEL staining, consistent with apoptosis.^6, 32, 33^ Retinal thinning has also been documented in living patients with diabetes.^34^ Importantly, this damage appears to be selective, with retinal ganglion cells and short-wavelength-sensitive (S) cone photoreceptors being particularly vulnerable.^35^ Similar findings have been reported in rodent models of streptozotocin-induced diabetes.^36^ Psychophysical studies have similarly demonstrated early functional deficits. In particular, abnormalities in Tritan color vision, initially attributed to changes in light scattering, have been shown to reflect neural dysfunction. These deficits can occur at an early stage of diabetes and may be partially improved by oxygen inhalation.^37^ Comparable abnormalities have also been demonstrated using other psychophysical measures, including assessments of photoreceptor sensitivity, color contrast sensitivity, and contrast sensitivity.^38–41^ Collectively, these findings support the view that diabetes induces early and selective neuronal and glial dysfunction in the retina, preceding or occurring alongside the more readily recognized vascular manifestations of diabetic retinopathy.

Recent studies have suggested that diabetic retinopathy is at least partly initiated by retinal photoreceptors exposed to metabolic stress.^18, 19^ However, the effect of elevated glucose on photoreceptor production of mediators capable of activating neighboring retinal cells has not been fully explored. Here we show that photoreceptors exposed to high glucose elicits increased secretion of VEGF, TNF-α, and IL-6. These mediators are associated with retinal inflammation and diabetic retinopathy progression, supporting the concept that photoreceptors can directly respond to hyperglycemia and initiate inflammatory responses from neighboring cells.

Retinal photoreceptors have traditionally received attention in diabetic retinopathy because of their high susceptibility to diabetes-related cell death and their oxygen consumption and potential contribution to retinal metabolic stress.^28, 29^ Our findings provide a complementary mechanism in which hyperglycemia directly alters photoreceptor secretory activity. Müller glial cells appear to amplify this photoreceptor-derived signal. After stimulation with products of photoreceptors exposed to high glucose, Müller glia produced approximately fourfold more VEGF than Müller cells exposed to high glucose alone. Müller cells also increased MMP-9 expression, secretion, and gelatinase activity. Together, these findings indicate that photoreceptor-derived signals can induce a sustained glial response involving both growth-factor production and extracellular-matrix remodeling.

Although MMP-9 has been implicated in the pathogenesis of diabetic retinopathy, its production by Müller cells raises the possibility of an additional mechanism linking Müller-cell dysfunction in diabetic retinopathy. Müller cells are anatomically positioned at the neurovascular interface and contribute to maintenance of the blood-retinal barrier.^42^ Previous studies have demonstrated that Müller cells can express and secrete MMP-9, particularly in response to inflammatory stimuli such as TNF-α.^43^ Thus, increased MMP-9 production by Müller cells under diabetic or inflammatory conditions could potentially promote extracellular-matrix remodeling in the perivascular microenvironment. MMP-9-mediated degradation of basement-membrane components and tightjunction-associated proteins could weaken endothelial barrier integrity, thereby increasing vascular permeability, contributing to retinal edema. Importantly, MMP-9 is produced by multiple retinal and infiltrating cell populations, and the relative contribution of Müller-cell-derived MMP-9 to the overall MMP-9 activity observed in diabetic retina remains to be determined.

## CONCLUSION

In summary, we provide evidence that photoreceptors exposed to high glucose produce increased levels of inflammatory and growth-regulatory mediators and that their secreted products activate Müller glial cells. High glucose photoreceptor-conditioned media increase Müller glial VEGF and MMP-9 production and MMP-9 enzymatic activity. These findings identify a direct photoreceptor-to-Müller glia activation pathway that may contribute to neuroglial inflammation in hyperglycemic retina and highlight photoreceptors as potential early cellular contributor to Neuroglial Inflammation.

## Acknowledgements

The authors thank Rong Yang and Gary McCollum for technical support.

## Author Contributions

MIU and IDH wrote the manuscript. JLN, JMP, YC, and DHZ performed the experiments, collected and analyzed the data. MIU, IDH and JSP conceived and supervised the project.

